# Social context modulates multibrain broadband dynamics and functional brain-to-brain coupling in the group of mice

**DOI:** 10.1101/2022.12.29.522176

**Authors:** Jeongyoon Lee, Damhyeon Kwak, Gwang Ung Lee, Chan Yeong Kim, Sang Hyun Park, Jee Hyun Choi, Sung Q. Lee, Han Kyoung Choe

## Abstract

Although mice are social animals, studies that explore the simultaneously recorded neural activities of multiple mice, especially in a social setting, are still lacking. In this study, we simultaneously recorded local field potential (LFP) signals in the dorsomedial prefrontal cortex (dmPFC) from up to four mice. The brain activities of the mice were measured in two contradicting conditions - freely interacting in a group or being individually separated. We found that social context and the locomotive states predominately modulate the entire LFP structure. Power spectral density (PSD) estimate and spectrogram of LFP signals showed a broadband modulation; lower frequency bands—delta (<4Hz), theta (4-7Hz), and alpha (8-12Hz) power were highly correlated to each other and anti-correlated with gamma and high gamma (>30Hz) power. We calculated the high-to-low-power ratio (HLR) and found that HLR was higher when the mice were in a group than were separated. The HLR was also higher when they were active—whether or not they were moving. The mice in the group showed higher HLR in any locomotive states. We then analyzed whether social context can be divided into sub-contexts. Notably, the aggregation of animals, called huddling, decreased social context-induced increase in HLR. Multibrain analyses of HLR indicated that the mice in the group displayed high cross-correlation to each other, indicating interbrain synchrony. Then we examined whether there is any directional relationship between HLR from pairs of mice. A majority of dyad selected within the group of mice showed unilateral precedence of HLR by Granger causality analysis, comprising a hierarchical social structure based on a directionality of influence. Overall, this study shows the importance of the social environment in brain dynamics and emphasizes the value of simultaneous multibrain recording for researching social behaviors and their neural correlates.

**One-sentence summary:** Coexistence modulates overall brain activities with unilateral causal relationship.

## Introduction

Social behaviors are built upon the brain activities which collectively occurs in the all participants. These brain activities are in turn subjected to be modulated by social context and social behavior. In social context, coordinated activities of brains, which is unpredictable from the sum of individual brain activities, emerges reportedly in a variety of species ranging from mouse, bat, monkey and human (Kingsbury et al., 2019; Redcay & Schilbach, 2019; Schippers et al., 2010; Tseng et al., 2018; Zhang & Yartsev, 2019). A variety of measures of brain activities, ranging from functional magnetic resonance imaging (fMRI), local field potential (LFP), calcium concentration at the single cell level, and firing rates of single units exhibits a coordinated and synchronous change among multiple animals in a group or dyad (Kim et al., 2020; Kingsbury et al., 2019; Montague et al., 2002; Rose et al., 2021; Zhang & Yartsev, 2019). On top of social coexistence, specific behavior patterns that are exclusively meaningful in social context, such as holding hands and ultrasound vocalization, can drive an additional level of synchronization in the brain activities (Goldstein et al., 2018; Rose et al., 2021). However, it has only begun to be explored how the combination of social context and behavioral status together shapes multibrain dynamics in all members of a group.

The representation of social information and the execution of social behaviors require an extensive brain network (Kingsbury & Hong, 2020; Redcay & Schilbach, 2019; Wei et al., 2021). In this network, the medial prefrontal cortex (mPFC) plays critical roles, ranging from social perception, social decision-making and mentalizing, in a phylogenetically conserved manner from rodents to humans (Amodio & Frith, 2006; Tremblay et al., 2017). The dorsomedial PFC (dmPFC) is involved in representing social information in a group of animals (Wang et al., 2011). The mPFC of a pair of mice shows correlated brain activities which can be utilized to predict the behavior and brain activities of the each other, and the frontal cortex of bats exhibited neural signals that are correlated and even reciprocally causal (Kingsbury et al., 2019; Rose et al., 2021; Zhang & Yartsev, 2019). Notably, artificial optogenetic stimulation of the dmPFC drives interaction of coherently stimulated pair, but not the animal stimulated out of phase (Yang et al., 2021), suggesting a causal role of dmPFC in social behavior.

The challenge of investigating a group of interacting animals exponentially grows as the number of subject animals increases. In spite of recent progress in multibrain recording technology (Rose et al., 2021; Shin et al., 2022), technical limitations, such as battery life packageable into head-mounted device and burden of meticulous behavioral annotation, prevented an experimental design long enough to entail a shift in social and behavioral states. We utilized huddling as easily identifiable social states. Huddling, a stable aggregation of animals, is both social and thermoadaptive behavior and is exclusively present in social conditions (Ito et al., 2019; Kojima & Alberts, 2011). In addition, we exploited head-mountable wireless edge-computing system capable of more than three-hour neural recording to understand multibrain dynamics of group of mice. Based on these, we addressed how brain dynamics are shaped by the interaction of locomotive states, group context, and social states, as well as the relationship of members in the group under task-free group conditions, by simultaneously recording the neural activity of the dmPFC and monitoring the behavior of animals.

## Results

### Broadband modulation of dmPFC LFP by locomotive state and group states

To address how group context shapes brain dynamics, we recorded the local field potential (LFP) of dmPFC of a set of the four mice once in group context, then followed by individual context (Fig. 1A). A group of four mice were placed in a single test box in the first experiment day (‘group’ session); each of the four mice was placed in four different boxes in the second experiment day (‘single’ session). During the entire behavioral session, we could obtain stable signal from the dmPFC transmitted from wireless head-mounted device (Fig. 1B). We examined the implant site *post hoc* and only included animals with the electrode located in the dmPFC (Fig. S1A). These recording sites mostly were located in the prelimbic area (PL) or anterior cingulate cortex (ACC) (Fig. S1B). Along with the LFP, we tracked and classified the locomotive states of each mouse into active, active-still, and inactive (Fig. 1C and D). The amplitude of the LFP appears to be related with the locomotive states in both recording conditions, with the higher amplitude of the LFP correlated with the inactive states. The time spent in either inactive, active-still, or active was not affected by social context (p>0.05) (Fig. S1C).

**Fig. 1.**
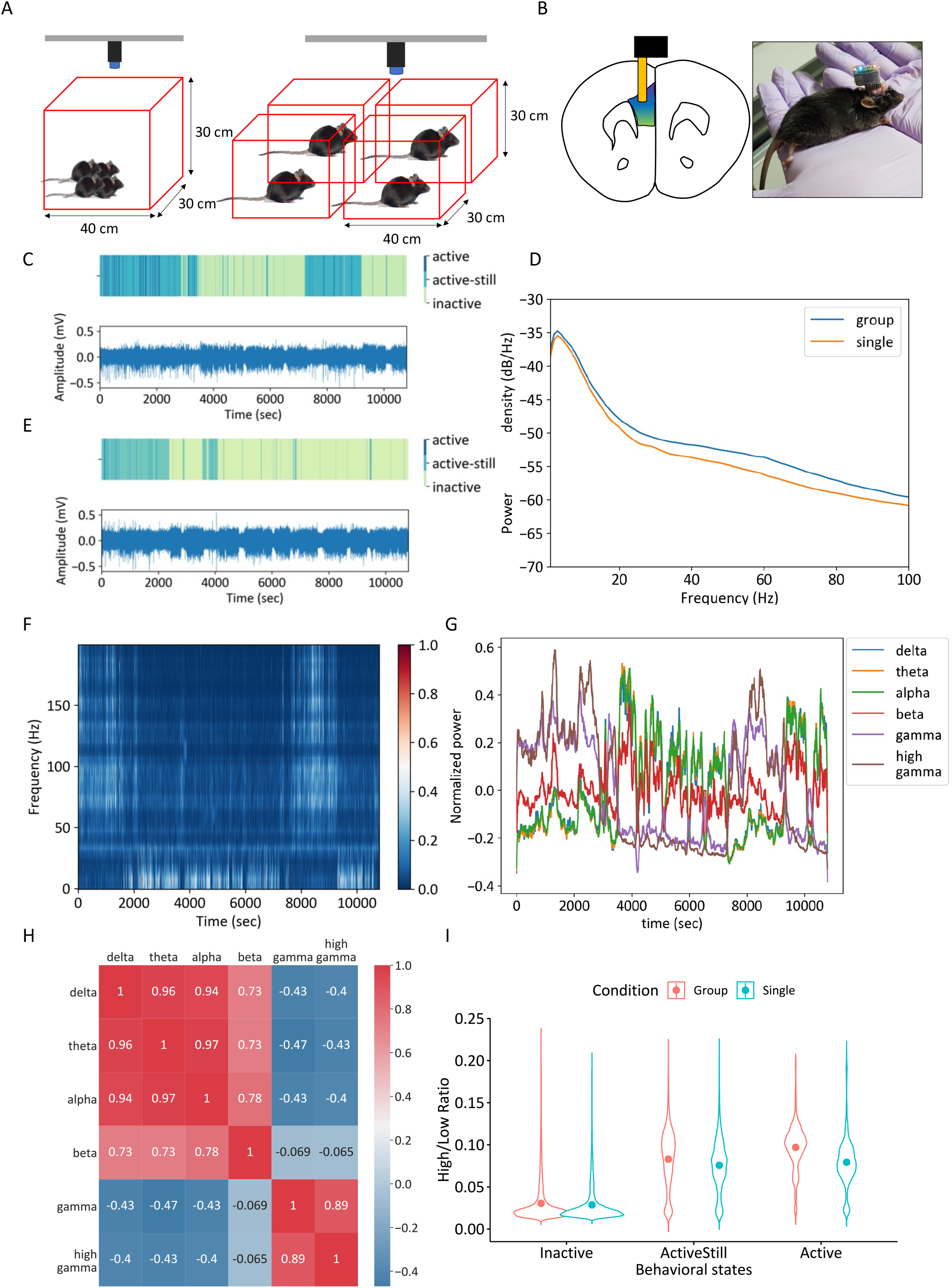
Behavior and dmPFC LFP patterns in group and single conditions. A: Group condition and B: Single condition. Mice have head-mounted wireless modules. C, E: Behavior and raw electrophysiology signals. D: Representative power spectral density (PSD) estimates of a mouse in two conditions. F: Representative normalized spectrogram of a mouse. G: Normalized power for each band calculated from F. H: Cross-correlation coefficient between different bands averaged from all mice and conditions. Delta-theta-alpha and gamma-high gamma band are highly correlated, while beta band has a weaker correlation.

To probe whether there are any frequency-specific signatures associated with group conditions, we first calculated the power spectral density (PSD) estimates of the LFP obtained during either group or single conditions (Fig. 1E). We found that averaged PSD signals differed by group and single conditions, with a stronger high frequency power in group conditions.

To better understand frequency characteristics of the dmPFC LFP, we obtained spectrograms of the LFP and calculated the average power of each brain wave band (Fig. 1F). The definitions of the band were as follows: delta (0-3Hz), theta (4-7Hz), alpha (8-12Hz), beta (12-30Hz), gamma (31-80Hz), and high gamma wave (81-150Hz). There was a broadband modulation in the spectrogram, with very high cross-correlations between delta, theta and alpha band powers (r > 0.9) and between gamma and high gamma band powers (r = 0.89). Delta/theta/alpha bands were also anti-correlated to gamma/high gamma bands (r=-0.47-0.4). These results are analogous with a previous observation made in bat frontal cortex, which is without the established LFP band distinction, that showed the opposing changes in the powers of high and low frequency band in active and resting conditions (Zhang & Yartsev, 2019). Additionally, beta bands had weaker correlation (r=0.73-0.78) to delta, theta and alpha bands and no correlation (r = -0.069-0.065) to gamma and high gamma bands (Fig. 1G and H). With these results, we defined ‘low’ band as 0-12 Hz (delta/theta/alpha) and ‘high’ band as 31-150 Hz (gamma/high gamma) for further analyses.

Based on this definition, we plotted the LFP power in low and high bands of each 1-second segment according to their locomotive states and social context (Fig. S1D). Scatter plots of both group and single conditions showed similar patterns. Inactive state forms a cluster that is distributed across varying degree of low bands power while maintaining the relatively weaker level of high bands power. In contrast, segments in both active-still and active states are located in another cluster that is scattered across varying degree of high bands power while maintaining relatively weaker power range of low bands. We then sought to analyze the effect of social context on LFP power in each locomotive state (Fig. 1I). Locomotive states were the major determinant of the dmPFC HLR, with stronger HLR observed in more mobile states. In each locomotive state, group conditions significantly increase the HLR compared to the same locomotive state in single conditions (locomotive state: p<0.05; group context: p<0.05 by repeated two-way ANOVA).

### Group-elicited increase in HLR is attenuated by huddling

In group context, mice display a variety of interactive behaviors. We hypothesized that a specific set of social behaviors may modulate brain activities in the dmPFC. For the sake of simplicity, we focused on huddling behavior as readily identifiable social states. Huddling behavior, defined as a stable physical aggregation of animals, serves a variety of social functions, such as predator evasion, as well as thermoadaptation (Harshaw et al., 2018; Kim et al., 2020). To address how the social states modulate the LFP in the dmPFC, we analyzed the HLR according to the huddling states along with locomotive states and group conditions. In grouped conditions, mice required slight latency to initiate huddling of a small size, then gradually forming larger ones (Fig. 2A). Once the full-sized huddling is formed, it maintained for more than thousands of seconds with occasional break up and re-huddling. The participation of individual mouse to huddling is analyzed in figure 2B. This pattern was consistent among all other groups that we analyzed (Fig. 2C). The huddling of size 4 occupied more than 60% of experimental session, while huddling of smaller size or non-huddled state filled the remaining portion. We quantitatively analyzed the distribution of locomotive state in either huddled or non-huddled state (Fig. 2D). Although the mice were dominantly inactive during huddling, they groomed themselves or huddled partner, presenting active-still state during huddling. They also switched the position inside the formed huddling presenting active state.

**Fig. 2.**
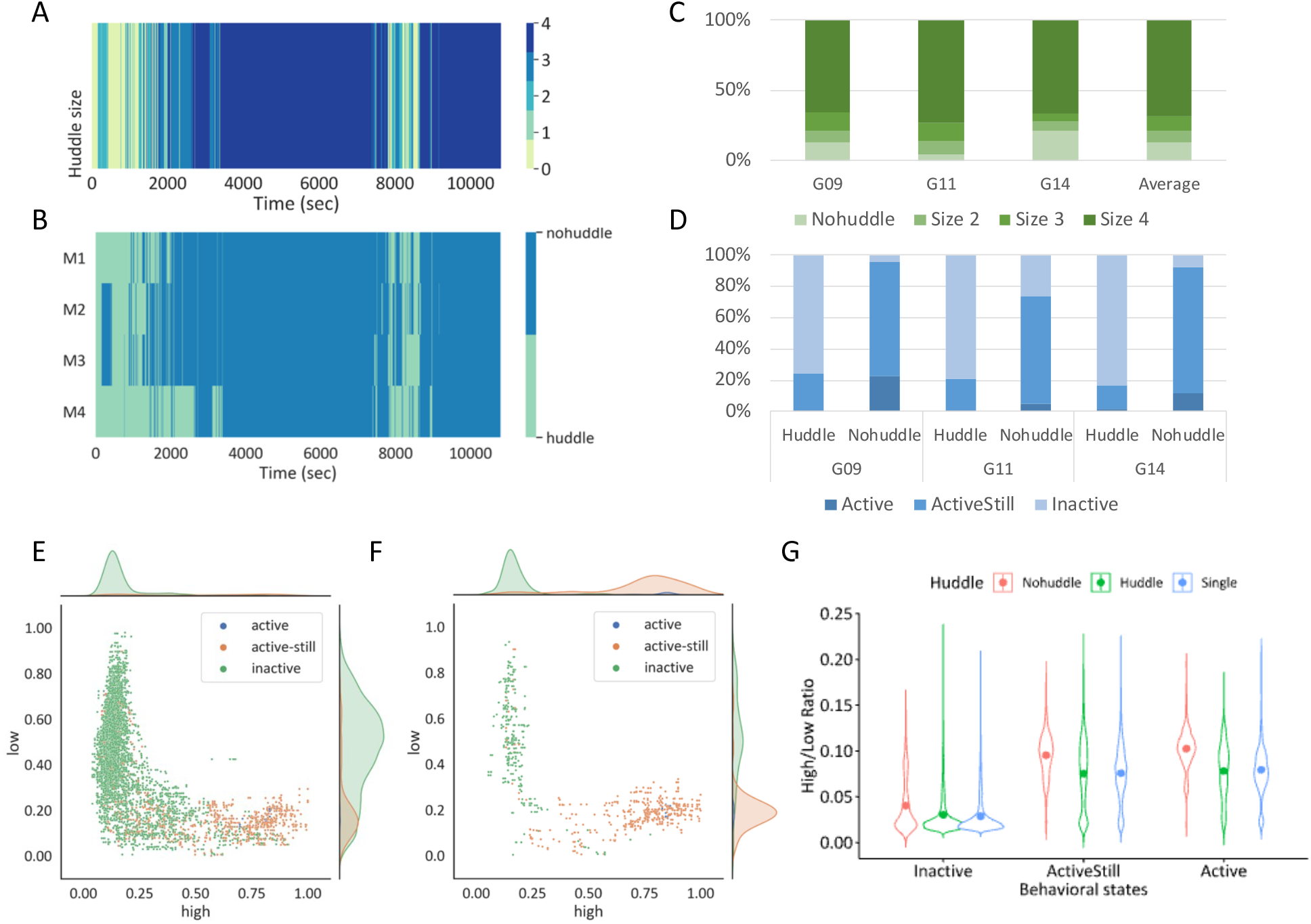
Modulation of HLR by huddling. A: Huddling state of grouped mice. Number indicates huddling size. B: Huddling state of each mouse in a group. C: % time spent for each huddling size. D: % time spent for each activeness state for huddling states. E, F: Scatterplot of normalized power for huddling (I) and non-huddling mouse (J). G: Average HLR by huddling states compared to single condition and activeness states.

We then combined behavioral results with extracellular recording data. We first plotted the LFP power of both high and low bands in huddling and non-huddled states (Fig. 2E and F, respectively). In both huddled and non-huddled states, the distribution of LFP power along locomotive states are segregated into two cluster, similarly to single or group conditions (Fig. S1D). Analysis of average HLR calculated based on each of huddling and locomotive states revealed a notable contribution of huddling in LFP modulation (Fig. 2G). In all locomotive state, the average HLR in huddled states are comparable to that of single conditions, while the HLR in non-huddled state was significantly upregulated compared to either single or huddled states. Considering the HLR in group conditions was higher than that in single conditions, non-huddled states account for the higher HLR in the group conditions. Our finding indicates that the huddling states segregate group conditions into two distinct social states, both in the view of behavior and brain activities.

### Interbrain correlation among the dmPFC activities in group conditions

We next set out to understand the interbrain relationship across the members of mice in group. We analyzed locomotive states of each mouse in single or group conditions (Fig. 3A and B, respectively). Mice under grouped conditions exhibited similar locomotive pattern with other co-housed members (Fig. 3A), in contrast to independent and out-of-sync activity onset and offset observed in single conditions (Fig. 3B). As all mice group that we examined spent a significant portion of time in huddling enriched with inactive state (Fig. 2C and D), huddling may provide a behavioral platform for interlinked inactive states across mice. In addition, non-huddled period of mice is predominantly occupied by active and active-still state (Figure 2D), providing another source of the similarity of locomotive state mediated by huddling. However, closely aligned onset and offset of locomotive states cannot be solely explained by the propensity of each locomotive states limited by huddled and non-huddled states.

**Fig. 3.**
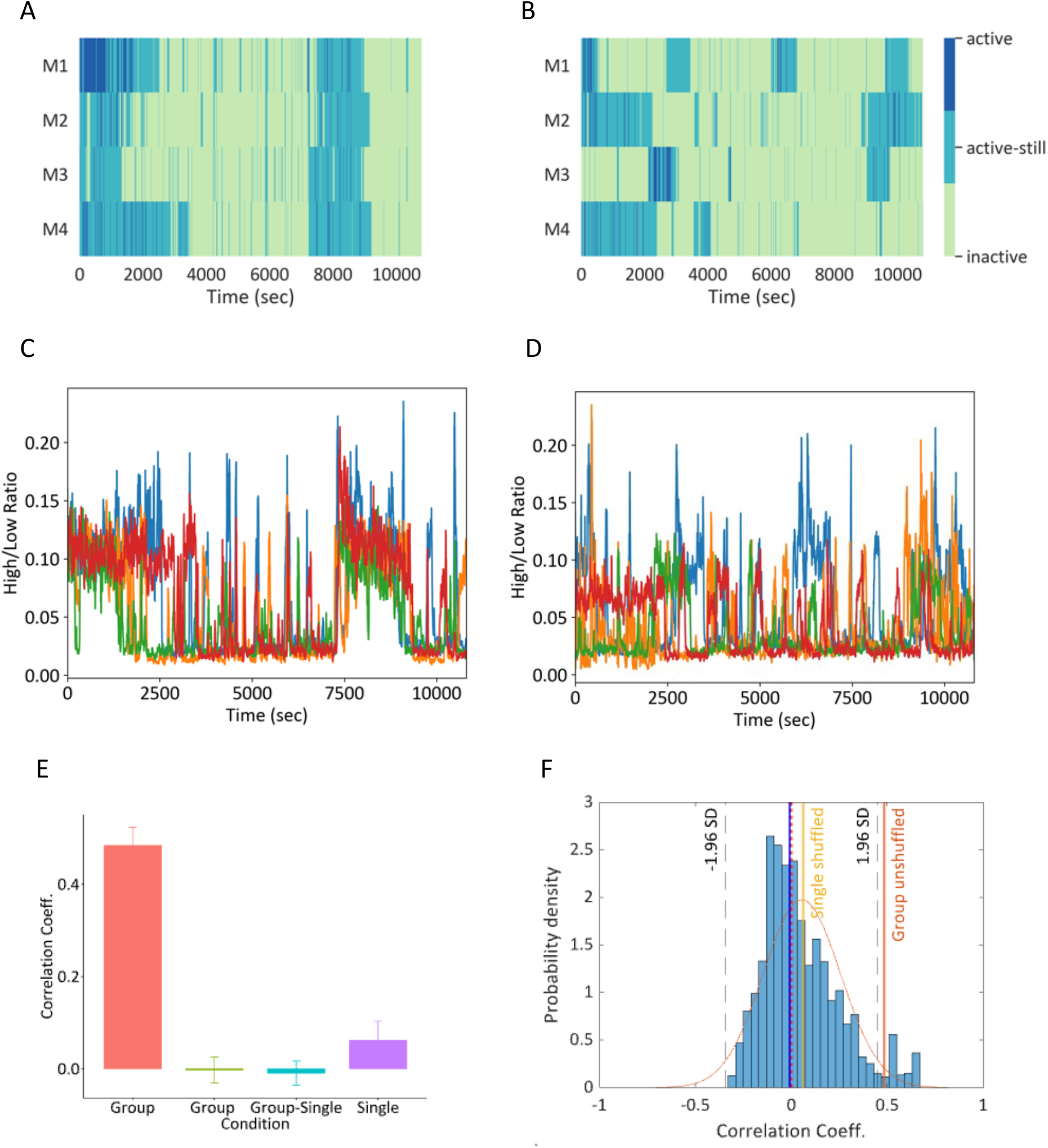
Interbrain correlation. A, B: Activeness status of mice in group (A) and single (B) condition. C, D: HLR of mice in group (A) and single (B) condition in time domain. E: Average interbrain cross-correlation of time domain HLR within group and for shuffled data. F: Bootstrapping of interbrain cross-correlation. Vertical line indicates mean values for each shuffling condition.

Then, we examined the brain activities of each member in mice group. The HLRs of each mouse plotted on the time scale were also correlated in the group conditions but not in single conditions (Fig. 3C and D), reflecting the similarity of locomotive states observed only under group conditions. To quantitatively analyze the correlation, we calculated the Pearson correlation coefficient for HLRs (Fig. 3E). The HLR correlation for a pair selected in the same group was higher than other shuffled pair: pairs selected across groups, pairs of one from group conditions and the other from single conditions, and pairs selected from single conditions. Full permutation tests of all possible pair in the dataset indicates that the average in-group correlation value of group context is at 2.5 percentile of the distribution, validating the statistical significance of the interbrain correlation exclusively observed in the group conditions (Fig. 2F). These results expand the interbrain correlation reported previously (Kingsbury et al., 2019; Zhang & Yartsev, 2019) to be persistent during hours of behavioral session in group of mice up to four.

### Unidirectional influence of mice group revealed by Granger causality

Given the high interbrain correlation of the HLRs in the mice in group conditions, we sought to find whether there is a unidirectional temporal organization of the brain activities across mice. Understanding unidirectional relationship between mice may provide us a novel way to investigate the group structure of mice. Group structure of mice that is determined by temporal relationship of brain activities may be all-to-all as depicted in Fig. 4A, hierarchical as depicted in Fig. 4B, or somewhere in between. We analyzed the profiles of HLR in a group of mice using Granger causality which can statistically examine whether one time-series data can usefully predict the other (Granger, 1969). Granger causality tests were conducted for all possible permutations for the mice in group (Fig. 4C-E). The HLR of certain mouse showed a significant temporal precedence over others. For instance, the HLR of mouse 1 in the group 9 preceded all other mice (mouse 1 precedes mouse 2, 3, and 4, p<0.05; mouse 2, 3, or 4 precedes mouse 1, p>0.05 by Granger causality test at lag=1), while the HLR of mouse 2 preceded mouse 3 and 4 but not mouse 1 (mouse 2 precedes mouse 3 and 4, p<0.05; mouse 2 precedes mouse 1, p>0.05)

**Fig. 4.**
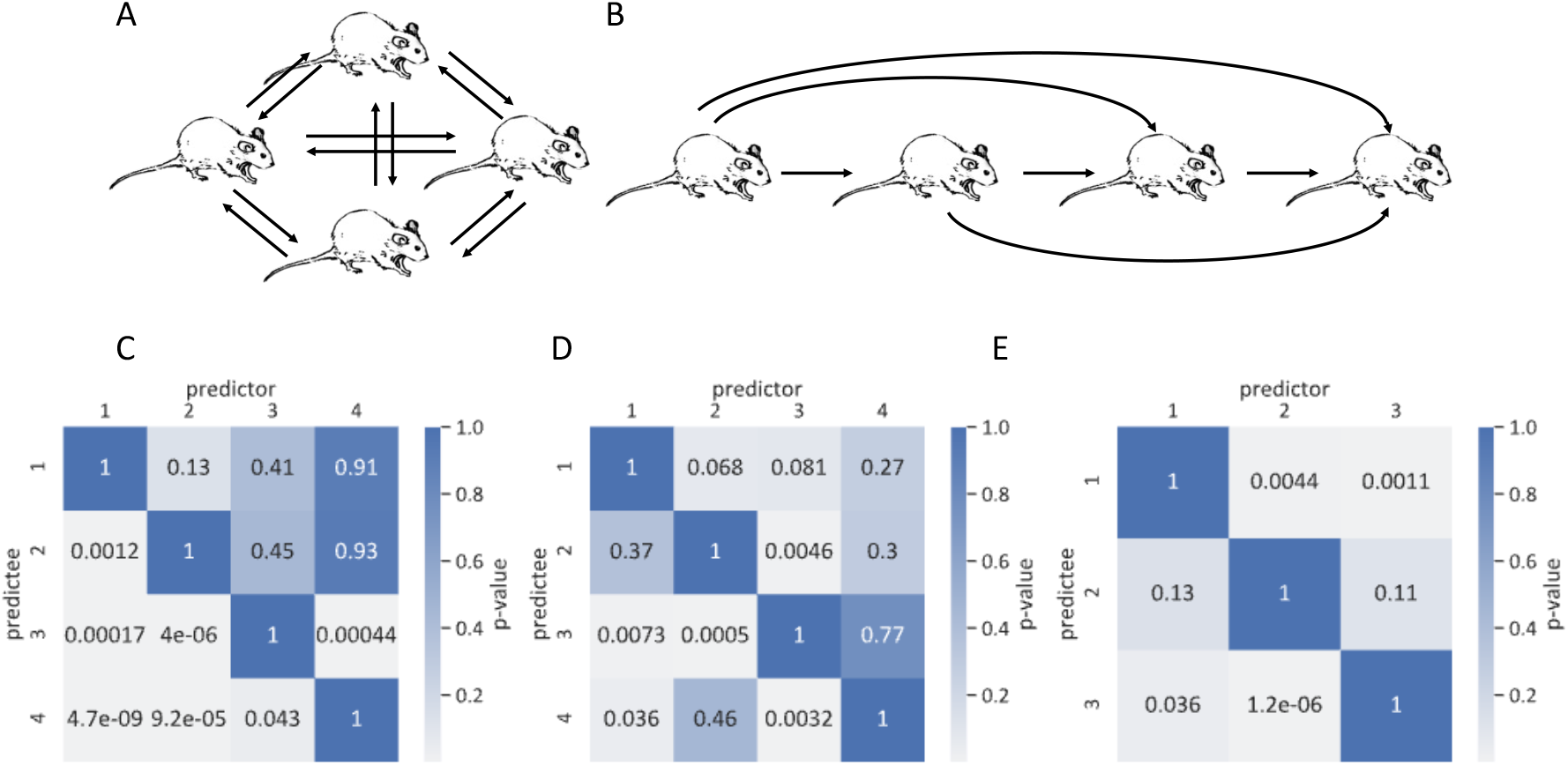
Directional influence between animals in group conditions. A, B: Possible scenarios of Granger causality. A: All mice granger cause bilaterally. B: Unilateral granger causality among mice implying temporal precedence in neural signal. C, D, E: Granger causality between HLR of mice in each group. Numbers inside squares indicate p-value where p<0.05 indicates granger causality.

(Figure 4C). We also found the similar organization of unilateral temporal precedence in other examined groups. These results indicate that interaction between each mouse in group is temporally directional. It also shows that unidirectional precedence of brain activities forms hierarchical structure among mice group in that the most preceding mice significantly forecasts the brain activities of all the other mice, and the second most preceding one forecasts the other two, but not the most preceding one, and the least preceding one forecasts none of the group members.

## Discussion

Here, we demonstrated that locomotive and social states are significant determinants of structure of dmPFC brain activities, as probed by the power ratio of the high frequency bands and low frequency bands (HLR). Notably, the modulatory effect of social context can be divided into two sub-contexts by the presence or absence of huddling. We then explored the interbrain relationship of neural activities. Significant correlation of HLR was detected between pairs in the same group. Analysis of temporal influence among members of the group revealed that there is unilateral influence to possibly build a hierarchy-like structure of the mouse group.

We noted the LFP modulation in the dmPFC by group context. The social situations may augment the change of the HLR due to an intensified or diversified locomotion that is driven by social interactions, or influence the HLR independent of the locomotion. To examine these hypotheses, we compared the HLRs in individual and social context with locomotive state-matching. The locomotion-matched HLR was increased in the social context compared to single conditions. In addition, the composition of locomotive states was not significantly different between group and single conditions. However, we cannot exclude the possibility that the difference may be, at least partly, due to an increased complexity of behavioral repertoire in the social situation. The similar logic can be applied to the change of HLR due to huddling states. When we compared the HLR in the corresponding locomotive states among single, non-huddled group, huddled group conditions, huddling decreased the social context-triggered increase in HLR back to those in single conditions. While all three locomotive states exist when the mice huddle, the detailed changes in behavioral microstates (which was not documented in the current study) may have affected the HLR decrease due to huddling. On other hand, there is a possible contribution to increase the mPFC activity in single conditions, such as heightened alertness due to isolation (Brandt et al., 2022). The neural basis of overall activity changes elicited by social sub-contexts should be further addressed in the future studies with a better resolution of behavioral annotation.

The current study shows the huddling behavior reverses the group-elicited increase in HLR of the dmPFC. The remaining question is: what would be contributing factor from huddling? The first factor to be considered is the temperature. It is well known that many animal species huddle for thermal regulation (Gilbert et al., 2009) and mice are not an exception (Batchelder et al., 1983). Yet, the relationship between the thermoregulatory behavior and dmPFC LFP modulation is not clearly identified. The other factor would be physical sensation, or touch itself. The slow and gentle touch is perceived as positive and affective (Mcglone et al., 2014), providing the basis of social affective touch. A close physical contact induced by huddling may provide such slow and gentle touch, which may contribute to social brain activities in the dmPFC.

A term “hyperscanning” refers to simultaneous multibrain recording. It has provided multitudes of the evidence for human interbrain synchrony with various modalities (Bilek et al., 2015; Kinreich et al., 2017). In animals, two groups have shown the interbrain synchrony with functional calcium imaging (Kingsbury et al., 2019) and with electrophysiology (Zhang & Yartsev, 2019) in two interacting mice and bats, respectively. Recent study has examined up to eight bats simultaneously and observed the interbrain correlation between them during their ultrasonic vocalizations (Rose et al., 2021). However, up to our knowledge, no study has reported the interbrain synchrony in mice more than two. In the current study, we have shown the interbrain synchrony in a larger group of mice up to four in a long-session trial. All four mice exhibited correlated patterns of the dmPFC HLR change in time when they were in a group, while there was no correlation of HLR when they were separated. The correlated neural activities were accompanied by the correlated behavioral states. It is currently unclear whether the neural activity of the dmPFC precedes the behavior or whether the behavior precedes the neural activity. Considering the recent studies that addressed sequential activation of the mPFC neurons along behavior sequence (Zhang et al., 2022), the temporal relationship of the neural activity and the behavior may not simply be preceding nor succeeding, rather require a further comprehensive understanding.

A group of mice forms a hierarchy-like structure (Zhou et al., 2018). The dominance-subordinate relationship of mice can be revealed by an arsenal of behavioral assays, for example, tube test. Recent progress indicated the factors determining social dominance and their neural substrate, while the communication between members comprising the hierarchy has not been much elaborated. We utilized the Granger causality analysis and demonstrated that the unilateral temporal precedence exists among the group of mice. In other words, the change of dmPFC activities of a certain mouse occurs earlier than that of other mice in the group. The naturally following question would be if this phenomenon is related to the social hierarchy of the mice group. There could be two possible scenarios: 1) the preceding mouse is the dominant one, so that it leads the group and others follow it; or oppositely, 2) the preceding mouse is the subordinate one where it has to act earlier than others for the better outcome/survival. The current study does not provide an answer to this question. It would be highly valuable to examine multibrain neural signal-behavior relationship with the behavioral paradigm probing social hierarchy to understand the relationship between the neural activity precedence and the social hierarchy.

The alteration of LFP observed in the wide frequency range of LFP may be due to either multiple changes of LFP powers in distinct frequency bands or overall shift of local neural network activity (Gao et al., 2017). Considering previous studies employed averaging of multiple session based on a reference point such as calling (Rose et al., 2021), pinpointing a reference timestamp to align neural activities in the current experimental conditions would help delineating the two possibilities. However, video tracking-based annotation of both locomotive and huddling status prevent a precise calling of behavioral states. Addition of sensor in the recording module or leading-edge machine vision techniques can help precise annotation of behavior that matches the temporal precision of neural activity analysis. Alternatively, the broadband modulation of LFP may be natural response of the brain during task-free and/or naturalistic environment.

In nature, house mice (*Mus musculus*) are social animals, and they tend to live together in a breeding unit called demes containing several mice (DeFries & McClearn, 1970; Singleton & Krebs, 2007). It is conceivable that multi-mice, multibrain study are necessary for understanding the natural social behavior and its neural signature of the mice. Ironically, the majority of previous studies regarding the social behavior of mice have recorded only single mouse, mostly due to technological constraints, with notable exceptions (Kim et al., 2020; Kingsbury et al., 2019; Shin et al., 2022). The current study demonstrated the modulation of dmPFC LFP by huddling behavior and the interbrain correlation between mice, which all require simultaneous multibrain recordings to be analyzed. To our knowledge, this is the first study that explored the social dynamics between mice more than two with simultaneous neural recording in a task-free environment. Furthermore, these multibrain studies can be also employed in the human studies. Recent human fMRI study recorded the neural activities from three humans simultaneously (Xie et al., 2020); these multibrain study will provide further understanding of our social behaviors and the corresponding neural activities.

## Acknowledgements

We thank undergraduate research interns of the ABC lab at DGIST. This work is supported by NRF-2020R1A2C4002156, NRF-2019M3C1B8090845, 22-RT-01, 22-CoE-BT-03, ETRI Grant (Collective Brain-Behavioral Modelling in Socially Interacting Group, 21YB1500).

## Materials and Methods

### 1. Experimental model and subject details

All procedures were approved by the Institutional Animal Care and Use Committee of Daegu Gyeongbuk Institute of Science and Technology (DGIST). All methods were carried out in accordance with the approved animal procedures and laboratory safety guidelines of DGIST. C57BL/6J mice were born and reared in standard mouse cages (16 * 36 * 12.5 cm^3^) with food and water available ad libitum. Mice were weaned at 3–4 weeks of age and housed together with sex-matched siblings with up to four animals per cage. Mice were maintained at a 12:12-h light/dark cycle at 22 ± 1°C. The number of total animals used in the study is 12 animals (C57BL/6J mice, 10-14weeks), and they consist of 3 experiment groups of four. Surgery performed on a 10-weeks mouse.

After surgery, mice had a recovery period in a single cage for a week. After recovery, they are regrouped with other mice that completed recovery. During regroup period, a dummy module was mounted to habituate the weight of the module. All mice participated in the Single Animal LFP Recording Session and the Group Animal LFP Recording Session. Each mouse adapted to the weight of the cage and module in the habituation session before participating in the experiment, and there was at least 2 days interval between the single and group experimental sessions.

### 2. Surgery and animal preparation

For all surgery, mice were anesthetized with intraperitoneal injection of ketamine-xylazine mixture solution (100mg/kg and 10mg/kg, respectively) and placed in stereotaxic apparatus. A tungsten wire electrode attached to a CBRAIN headstage was inserted in dmPFC (AP=+1.70, DV=+1.65 and ML=+0.3). The tungsten wire electrode was coated with Di/I for locating the electrode in the brain prior to the surgery. The electrode was made of tungsten wire (114.3 μm thickness, PFA-Coated Tungsten Wire 796000, AM Systems) and was soldered with a thin wire connected to a 2×8 pinhead socket. Exposed metal was coated with resin (The Bondic Starter Pack, Bondic) for insulation. The impedance of the tungsten wire electrode was measured in vivo at 30-50kOhm at 1 kHz through a CBRAIN program. Two bone screws were placed into the skull on cerebellum and used as reference and ground electrodes. Inserted electrodes were fixed with a light cure adhesive (OptiBond All-in-One, Kerr) and that covered by dental cement. The mice went through the recovery period for one week after surgery. After the recovery, mice were mounted with dummy modules during anesthesia with isoflurane (Isoflurane USP, GNH India) a week before the experimental session for habituation.

The mice were colored by marker for identification three days before the experimental session. The mice were anesthetized with intraperitoneal injection of ketamine-xylazine mixture solution (100mg/kg and 10mg/kg, respectively). The bleaching agent was mixed and applied it to the back (neck side) of the anesthetized mouse with a size of 2.5cm-diameter circle. The bleaching agent was rinsed with water after 15 minutes. The fur was dried with an infrared heater and dyed with the marker.

### 3. Electrophysiological recording

Simultaneous wireless local field potential (LFP) recordings were performed with an implanted custom module called CBRAIN. The CBRAIN module used in this study was described previously (Kim et al., 2020). Briefly, it is a complete mobile edge computing solution consisting of an amplifier (RHD2216, Intan Technologies), telemetry based on a Bluetooth SoC (nRF52832, Nordic Semiconductor), a Cortex-M4 microprocessor (embedded in the Bluetooth SoC), power supply and LEDs in a small headstage with a weight of 2.6g, inclusive of a 2.0g lithium polymer battery. A custom GUI software (CBRAIN Studio) written in MATLAB was used for the data acquisition. The sampling frequency was set at 1024Hz. The data in a hexadecimal, space-delimited text format was transmitted via Bluetooth Enhanced ShockBurst broadcasting scheme to the recording computer. The data were segmented by a 30-minute interval and each segment was saved onto a separate text file.

### 4. Experimental sessions

We performed the group condition experiments first followed by the single condition experiment with a gap of a day. First, mice went through the habituation session in an experimental cage (Red acrylic, 40*30*30 cm^3^) for twelve hours. For group conditions, a cohort of four mice was introduced into one experimental cage together; for single conditions, each mouse was introduced into each of four cages individually.

At the Zeitgeber time (ZT) 0 which corresponds to the light-on time, the dummy module was replaced with the CBRAIN module under isoflurane anesthesia. The subject mice were returned to the experimental cage. Recording began 30 minutes after the mice were recovered from anesthesia. The camera () was installed at a height of 80 cm above the cage. We recorded the behavior of the subjects while they freely behave without a specific task for three hours. After the experiment, the mice were returned to their home cage.

### 5. Histology and confocal microscopy

After all experiment sessions, mice were intracardially perfused with 4% paraformaldehyde and post-fixed at 4 °C overnight. To inspect the injection sites, coronal sections (50μm thickness) were obtained using a Leica VT1000 S vibratome (Leica, Wetzlar, Germany). A sample of brains were obtained and sliced by cryosection. The brains were immersed in 30% sucrose in PBS and dehydrated for 3 d at 4 °C. The brains were embedded in optimal cutting temperature (OCT) compound (Scigen, Paramount, CA, USA) and frozen at − 80 °C. Coronal sections (40-μm thick) were obtained using a Leica CM3050 S cryostat (Leica). The prepared slices were labeled using 4’,6-diamidino-2-phenylindole (DAPI) and images were captured using a Nikon C2+ confocal microscope system (Nikon, Tokyo, Japan).

### 6. Behavior data analysis

Mouse behaviors were annotated manually by inspecting the video recording for each 1-second segment. ‘Active’ state is defined as the mouse changing its location. ‘Active-still’ state was defined as the mouse moving its head without displacement. ‘Inactive’ state was defined as the mouse showing no movement. For the group condition, huddling status was additionally annotated from the video recording. Huddling was defined as the mice touching their body longer than 10 seconds. The number of huddling mice and the participation status of the mice were recorded. Sleep was defined as consecutive immobility for two minutes (ref.). The annotated behavioral pattern was visualized by the Seaborn (Version) package in Python 3.9.

### 7. Electrophysiological data analysis

#### a. Data preprocessing

The segmented hexadecimal data obtained from CBRAIN were converted into the decimal format and concatenated on a custom MATLAB 2021b software. Data from relevant channels were saved onto the MAT format file for further analysis. The MAT files were loaded on Python using the SciPy 1.7.1 or mat73 0.50 package. A low-pass filter with cutoff frequency at 200Hz, and a notch filters with frequencies at 60Hz and 120Hz were applied to the data using the SciPy package.

#### b. Power spectral density and spectrogram analysis

To obtain the average Power spectral density (PSD) estimates, the signal was first segmented by 1-second interval on Python. Welch PSD function with a hamming window from SciPy package as applied on each interval and averaged. The same procedure was repeated with the signals classified by concurrent behavioral data. Spectrogram of the data were calculated by the spectrogram function in the SciPy package.

The spectrogram was downsampled to have 1-second time resolution to match the behavioral data and normalized by min-max scaling using scikit-learn 1.0 package in Python if necessary. The average power for each brain wave bands were calculated for each 1-second segment. The definition of the bands was as follows: delta wave (0-3Hz), theta wave (4-7Hz), alpha wave (8-12Hz), beta wave (12-30Hz), gamma wave (31-80Hz), and high gamma wave (81-150Hz). Time series cross-correlation coefficients were calculated between all brain wave bands using the statsmodels 0.13.2 package in Python. The average of correlation coefficients obtained from each LFP recording was plotted on the heatmap. Based on the correlation analysis, we defined ‘low’ and as delta/theta/alpha (0-12Hz) and ‘high’ band as gamma/high gamma (31-150Hz) bands. The High/Low Ratio (HLR) was calculated by dividing average high band (31-150Hz) power by average low band (0-12Hz) power for each 1-second segment from unnormalized spectrogram. Scatterplots of average high and low band power were produced by jointplot function from the Seaborn 0.11.1 package in Python.

#### c. Interbrain correlation and Granger causality

The HLRs from each group and condition were used. The time-series cross-correlation coefficients were calculated in MATLAB for different combination of HLRs. ‘Group’ combination was HLRs from single group (e.g. only group 9) and from group condition; ‘Group shuffled’ combination was HLRs from different groups (e.g. group 9 and 11) and from group condition; ‘Group-Single shuffled’ combination was HLRs from single group but from both group and single condition; and ‘Single shuffled’ combination was HLRs from single group and from single condition. The latter three combinations served as negative controls. For permutation analysis, all HLRs from all groups and conditions were used. Two random HLRs were obtained from the pool to calculate correlation coefficients, and this procedure was repeated for 10,000 times.

Granger causality (GC) analysis was performed using grangercausalitytests function from the statsmodel package. Only HLRs from group condition were used. For each group, GC was calculated for all possible permutations of mice (e.g. 1->2, 1->3, 1->4, 2->1, 2->3, 2->4, 3->1, 3->2, 3->4, 4->1, 4->2, 4->3). GC p-values at lag=1 were plotted on heatmap for each group.

### 8. Statistical analysis

Statistical analysis was performed using R 4.2.1, SPSS 28, GraphPad Prism 8.01 and MATLAB R2021b. One-way repeated-measures ANOVA was used to compare the HLRs for the different behavioral states and conditions. Two-way ANOVA was used to compare distribution of locomotive states in different experimental conditions. One-way ANOVA was used to compare the correlation coefficient of different combinations of pairs.

## References

Amodio, D. M., & Frith, C. D. (2006). Meeting of minds: the medial frontal cortex and social cognition. Nature Reviews Neuroscience, 7(4), 268–277. https://doi.org/10.1038/nrn1884

Batchelder, P., Kinney, R. O., Demlow, L., & Lynch, C. B. (1983). Effects of temperature and social interactions on huddling behavior in Mus musculus. Physiol Behav, 31(1), 97–102. https://doi.org/10.1016/0031-9384(83)90102-6

Bilek, E., Ruf, M., Schäfer, A., Akdeniz, C., Calhoun, V. D., Schmahl, C., Demanuele, C., Tost, H., Kirsch, P., & Meyer-Lindenberg, A. (2015). Information flow between interacting human brains: Identification, validation, and relationship to social expertise. Proceedings of the National Academy of Sciences, 112(16), 5207–5212. https://doi.org/10.1073/pnas.1421831112

Brandt, L., Liu, S., Heim, C., & Heinz, A. (2022). The effects of social isolation stress and discrimination on mental health. Translational Psychiatry, 12(1). https://doi.org/10.1038/s41398-022-02178-4

DeFries, J. C., & McClearn, G. E. (1970). Social Dominance and Darwinian Fitness in the Laboratory Mouse. The American Naturalist, 104(938), 408–411. https://doi.org/10.1086/282675

Gao, R., Peterson, E. J., & Voytek, B. (2017). Inferring synaptic excitation/inhibition balance from field potentials. NeuroImage, 158, 70–78. https://doi.org/10.1016/j.neuroimage.2017.06.078

Gilbert, C., Mccafferty, D., Le Maho, Y., Martrette, J.-M., Giroud, S., Blanc, S., & Ancel, A. (2009). One for all and all for one: the energetic benefits of huddling in endotherms. Biological Reviews, no-no. https://doi.org/10.1111/j.1469-185x.2009.00115.x

Goldstein, P., Weissman-Fogel, I., Dumas, G., & Shamay-Tsoory, S. G. (2018). Brain-to-brain coupling during handholding is associated with pain reduction. Proceedings of the National Academy of Sciences, 115(11), E2528–E2537. https://doi.org/10.1073/pnas.1703643115

Granger, C. W. J. (1969). Investigating Causal Relations by Econometric Models and Cross-spectral Methods. Econometrica, 37(3). https://doi.org/10.2307/1912791

Harshaw, C., Leffel, J. K., & Alberts, J. R. (2018). Oxytocin and the warm outer glow: Thermoregulatory deficits cause huddling abnormalities in oxytocin-deficient mouse pups. Hormones and Behavior, 98, 145–158. https://doi.org/10.1016/j.yhbeh.2017.12.007

Ito, W., Huang, H., Brayman, V., & Morozov, A. (2019). Impaired social contacts with familiar anesthetized conspecific in CA3-restricted brain-derived neurotrophic factor knockout mice. Genes, Brain and Behavior, 18(1), e12513. https://doi.org/10.1111/gbb.12513

Kim, J., Kim, C., Han, H. B., Cho, C. J., Yeom, W., Lee, S. Q., & Choi, J. H. (2020). A bird’s-eye view of brain activity in socially interacting mice through mobile edge computing (MEC). Sci Adv, 6(49). https://doi.org/10.1126/sciadv.abb9841

Kingsbury, L., & Hong, W. (2020). A Multi-Brain Framework for Social Interaction. Trends Neurosci, 43(9), 651–666. https://doi.org/10.1016/j.tins.2020.06.008

Kingsbury, L., Huang, S., Wang, J., Gu, K., Golshani, P., Wu, Y. E., & Hong, W. (2019). Correlated Neural Activity and Encoding of Behavior across Brains of Socially Interacting Animals. Cell, 178(2), 429-446.e416. https://doi.org/10.1016/j.cell.2019.05.022

Kinreich, S., Djalovski, A., Kraus, L., Louzoun, Y., & Feldman, R. (2017). Brain-to-Brain Synchrony during Naturalistic Social Interactions. Scientific Reports, 7(1). https://doi.org/10.1038/s41598-017-17339-5

Kojima, S., & Alberts, J. R. (2011). Oxytocin mediates the acquisition of filial, odor-guided huddling for maternally-associated odor in preweanling rats. Hormones and Behavior, 60(5), 549–558. https://doi.org/10.1016/j.yhbeh.2011.08.003

Mcglone, F., Wessberg, J., & Olausson, H. (2014). Discriminative and Affective Touch: Sensing and Feeling. Neuron, 82(4), 737–755. https://doi.org/10.1016/j.neuron.2014.05.001

Montague, P. R., Berns, G. S., Cohen, J. D., McClure, S. M., Pagnoni, G., Dhamala, M., Wiest, M. C., Karpov, I., King, R. D., Apple, N., & Fisher, R. E. (2002). Hyperscanning: simultaneous fMRI during linked social interactions. NeuroImage, 16(4), 1159–1164. https://doi.org/10.1006/nimg.2002.1150

Redcay, E., & Schilbach, L. (2019). Using second-person neuroscience to elucidate the mechanisms of social interaction. Nature Reviews Neuroscience, 20(8), 495–505. https://doi.org/10.1038/s41583-019-0179-4

Rose, M. C., Styr, B., Schmid, T. A., Elie, J. E., & Yartsev, M. M. (2021). Cortical representation of group social communication in bats. Science, 374(6566), eaba9584. https://doi.org/10.1126/science.aba9584

Schippers, M. B., Roebroeck, A., Renken, R., Nanetti, L., & Keysers, C. (2010). Mapping the information flow from one brain to another during gestural communication. Proceedings of the National Academy of Sciences, 107(20), 9388–9393. https://doi.org/10.1073/pnas.1001791107

Shin, H., Byun, J., Roh, D., Choi, N., Shin, H. S., & Cho, I. J. (2022). Interference-free, lightweight wireless neural probe system for investigating brain activity during natural competition. Biosens Bioelectron, 195, 113665. https://doi.org/10.1016/j.bios.2021.113665

Singleton, G., & Krebs, C. (2007). The Secret World of Wild Mice. In The Mouse in Biomedical Research (pp. 25–51). https://doi.org/10.1016/b978-012369454-6/50015-7

Tremblay, S., Sharika, K. M., & Platt, M. L. (2017). Social Decision-Making and the Brain: A Comparative Perspective. Trends in Cognitive Sciences, 21(4), 265–276. https://doi.org/10.1016/j.tics.2017.01.007

Tseng, P.-H., Rajangam, S., Lehew, G., Lebedev, M. A., & Nicolelis, M. A. L. (2018). Interbrain cortical synchronization encodes multiple aspects of social interactions in monkey pairs. Scientific Reports, 8(1). https://doi.org/10.1038/s41598-018-22679-x

Wang, F., Zhu, J., Zhu, H., Zhang, Q., Lin, Z., & Hu, H. (2011). Bidirectional control of social hierarchy by synaptic efficacy in medial prefrontal cortex. Science, 334(6056), 693–697. https://doi.org/10.1126/science.1209951

Wei, D., Talwar, V., & Lin, D. (2021). Neural circuits of social behaviors: Innate yet flexible. Neuron, 109(10), 1600–1620. https://doi.org/10.1016/j.neuron.2021.02.012

Xie, H., Karipidis, I. I., Howell, A., Schreier, M., Sheau, K. E., Manchanda, M. K., Ayub, R., Glover, G. H., Jung, M., Reiss, A. L., & Saggar, M. (2020). Finding the neural correlates of collaboration using a three-person fMRI hyperscanning paradigm. Proceedings of the National Academy of Sciences, 117(37), 23066–23072. https://doi.org/10.1073/pnas.1917407117

Yang, Y., Wu, M., Vázquez-Guardado, A., Wegener, A. J., Grajales-Reyes, J. G., Deng, Y., Wang, T., Avila, R., Moreno, J. A., Minkowicz, S., Dumrongprechachan, V., Lee, J., Zhang, S., Legaria, A. A., Ma, Y., Mehta, S., Franklin, D., Hartman, L., Bai, W., … Rogers, J. A. (2021). Wireless multilateral devices for optogenetic studies of individual and social behaviors. Nature Neuroscience, 24(7), 1035–1045. https://doi.org/10.1038/s41593-021-00849-x

Zhang, W., & Yartsev, M. M. (2019). Correlated Neural Activity across the Brains of Socially Interacting Bats. Cell, 178(2), 413-428.e422. https://doi.org/10.1016/j.cell.2019.05.023

Zhang, Y., Denman, A. J., Liang, B., Werner, C. T., Beacher, N. J., Chen, R., Li, Y., Shaham, Y., Barbera, G., & Lin, D. T. (2022). Detailed mapping of behavior reveals the formation of prelimbic neural ensembles across operant learning. Neuron, 110(4), 674–685 e676. https://doi.org/10.1016/j.neuron.2021.11.022

Zhou, T., Sandi, C., & Hu, H. (2018). Advances in understanding neural mechanisms of social dominance. Curr Opin Neurobiol, 49, 99–107. https://doi.org/10.1016/j.conb.2018.01.006

