## Supplementary figure 1 for "Social context modulates multibrain broadband dynamics and functional brain-to-brain coupling in the group of mice"

Figure S1

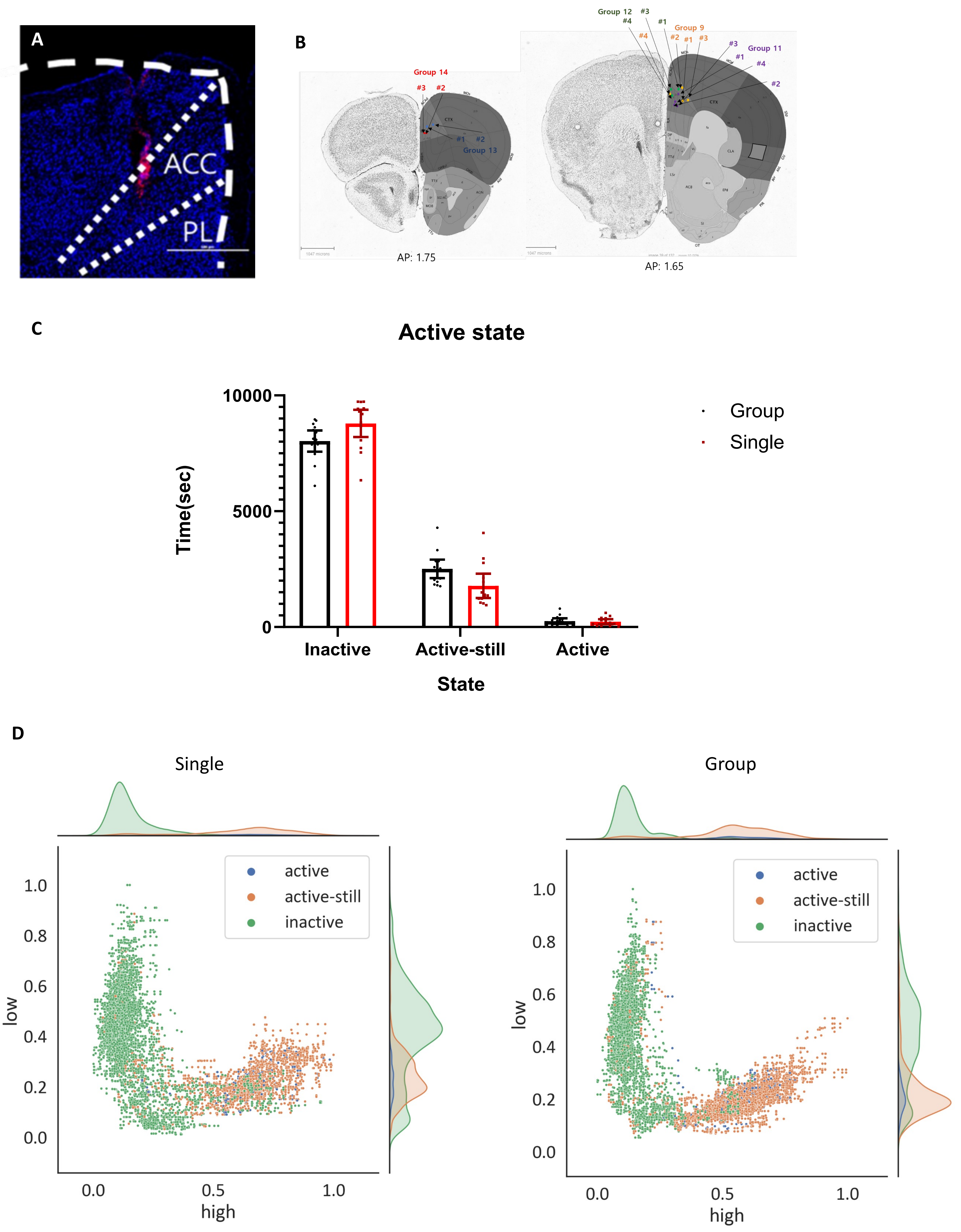


Figure S1. A. Representative brain section showing recording electrode implant site. Blue: DAPI, Red: DiI. Scale bar: 500 µm. B. Recording sites registered on the reference brain atlas. C. Propensity of locomotive states in group and single conditions. $P>0.05$ by two-way ANOVA. D. Scatter plots showing the high bands power and the low bands power of locomotive states in single (left panel) and group conditions (right panel).
